# Disentangling unspecific and specific transgenerational immune priming components in host-parasite interactions

**DOI:** 10.1101/429498

**Authors:** Frida Ben-Ami, Christian Orlic, Roland R. Regoes

## Abstract

Exposure to a pathogen primes many organisms to respond faster or more efficiently to subsequent exposures. Such priming can be unspecific or specific, and has been found to extend across generations. Disentangling and quantifying specific and unspecific effects is essential for understanding the genetic epidemiology of a system. By combining a large infection experiment and mathematical modeling, we disentangle different transgenerational effects in the crustacean model *Daphnia magna* exposed to different strains of the bacterial parasite *Pasteuria ramosa*. In the experiments, we exposed hosts to a high-dose of one of three parasite strains, and subsequently challenged their offspring with multiple doses of the same or a different strain, i. e. homologously or heterogously. We find that exposure to *Pasteuria* decreases the susceptibility of a host’s offspring by approximately 50%. This transgenerational protection is not larger for homologous than for heterologous parasite challenges. Our work represents an important contribution not only to the analysis of immune priming in ecological systems, but also to the experimental assessment of vaccines. We present for the first time an inference framework to investigate specific and unspecific effects of immune priming on the susceptibility distribution of hosts — effects that are central to understanding immunity and the effect of vaccines.

**Author summary:** Immune memory is a feature of immune systems that forms the basis of vaccination. In opposition to textbook accounts, the ability to specifically remember previous exposures has been found to extend to invertebrates and shown to be able to be passed on from mother to offspring, i. e. to be *transgenerational*. In this paper, we investigate the extent of this specificity in unprecedented detail in water fleas. We exposed water flea mothers to different strains of a bacterial pathogen and challenged their offspring with a wide range of doses of a strain that were either identical to (homologous) or different from (heterologous) the strain, to which the mother had been exposed. We find that, while exposure of the mother reduces the susceptibility of the offspring, this effect is not specific. This work outlines the limits of specific transgenerational immune memory in this invertebrate system.

## 1 Introduction

Transgenerational effects occur when the phenotype of the parent affects the phenotype of its offspring in addition to the direct effects of the genes contributed by the parent (Räsänen and Kruuk, 2007, Badyaev and Uller, 2009, Wolf and Wade, 2009). These effects are ubiquitous in nature and have been documented in a wide range of traits and taxa (Bernardo, 1996, Rossiter, 1996, Mousseau and Fox, 1998, Hereford and Moriuchi, 2005, Beckerman et al., 2006).

Among the most widely-studied transgenerational effects are those involving the transfer of immunity or increased parasite resistance from parents to offspring, commonly found in vertebrates (Grindstaff et al., 2003, Hasselquist and Nilsson, 2009), but also in invertebrates (Little and Kraaijeveld, 2004, Milutinović and Kurtz, 2016, Pigeault et al., 2016). The latter is particularly intriguing, because until the beginning of this millenium invertebrates were thought to have only innate immunity, which would render them naive whenever they encounter parasites (see, for example, Hoffmann and Reichhart (2002)). The potential of innate immune systems to specifically remember previous exposures to pathogens was first supported by phenomenological evidence (reviewed by Kurtz (2005), Schmid-Hempel (2005b,a)). In particular, it has been shown that invertebrate hosts can be primed against specific parasite species and strains (Kurtz and Franz, 2003, Sadd and Schmid-Hempel, 2006, Pham et al., 2007, Roth et al., 2009) or across life stages (Tate and Rudolf, 2012). There is also growing evidence that the immune system of invertebrates shares several homologies with that of vertebrates (Schmid-Hempel, 2005b, Watson et al., 2005, Kurtz and Armitage, 2006), although it has been argued that immune memory in invertebrates may be mediated by yet unidentified mechanisms that will not be found by looking for homologies (Little et al., 2005). More recently, the interest in potential innate immune memory is flaring up, and new molecular mechanisms are being elucidated in invertebrates (Barribeau et al., 2016, Tate et al., 2017), and even in vertebrates (Netea et al., 2016).

Immune priming can be unspecific or specific. Specificity here defines the degree to which a primed immune response is able to discriminate among different parasite strains, species or taxa (e.g., Gram-positive bacteria; Milutinović and Kurtz (2016)). While unspecific immune priming is important for eliciting a general response against a variety of parasites, specific immune priming can provide a targeted, and often more effective and long-lasting protection against reinfections. It is thus crucial to disentangle between unspecific and specific immune priming, in order to understand which of the two is responsible for an observed immune response.

Studies of transgenerational effects on disease typically subject the parental environment to food stress, for example, shortage of food or food of lower quality (Ben-Ami et al., 2010, Boots and Roberts, 2012), crowding (Michel et al., 2016) or challenge them with live, weakened or heat-killed parasites (Roth et al., 2010, Hernández López et al., 2014, McNamara et al., 2014). Thereafter, a variety of traits of the offspring generation are recorded, such as parasite infectivity and offspring fecundity, resistance, immunity and mortality (Hall and Ebert, 2012, Schlotz et al., 2013, Pigeault et al., 2015). Of the studies in which the parent generation was challenged with parasites, the vast majority investigated unspecific immune priming, i.e., exposing the mother generation to a parasite results in reduced susceptibility of its offspring, but this reduction in susceptibility was unaffected by the parasite strains or species used to infect the parent vs. offspring generation (Huang and Song, 1999, Sadd et al., 2005, Moret, 2006, Roth et al., 2010, Tidbury et al., 2011, Hernández López et al., 2014, McNamara et al., 2014). Only a handful of studies involving invertebrates found evidence for specific transgenerational immune priming. For example, Norouzitallab et al. (2016) showed the occurrence of specific immune memory in the brine shrimp *Artemia franciscana*, as manifested by increased resistance of the progenies of Vibrio-exposed ancestors towards a homologous bacterial strain, rather than to a heterologous strain. Little et al. (2003) obtained similar results in the crustacean *Daphnia magna* by exposing mothers to one parasite strain and testing their offsprings susceptibility to the same and different parasite strain.

Since specific immune priming can play an important role in host-parasite interactions at the population level, we combined an experiment and mathematical model to disentangle transgenerational effects of unspecific and specific immune priming in *Daphnia magna* challenged with different strains of its bacterial parasite *Pasteuria ramosa*. Instead of exposing host individuals to a single challenge dose of the pathogen, as is done in most studies, we use seven challenge doses ranging over more than 5 orders of magnitude. We chose multiple challenge doses because this allows us to study not just the average susceptibility but the entire distribution of susceptibilities in the host population. In particular, this approach can identify if there are subpopulations of hosts that respond to priming differently, rather than a uniformous priming response in every host individual — similar in principle to the concepts of all-or-none versus leaky vaccine efficacies conceived in the context of vaccine trials Struchiner et al. (1990), Halloran et al. (1992, 1996), Longini and Halloran (1996), Halloran et al. (2010).

## 2 Results

### 2.1 Dose dependence of infection rates

To determine the existence and extent of these various forms of transgenerational immune priming, we conducted experiments with *Daphnia magna* and three strains of its parasite *Pasteuria ramosa*, P1, P2, and P5. We first exposed genetically identical *Daphnia* to a high dose of one of the three parasites strains. As a control, some *Daphnia* were not exposed to any parasite strain. All unexposed control animals remained uninfected throughout the experiment. Overall, this resulted in four treatment groups.

The exposed and control *Daphnia* subsequently produced offspring. Offspring from the four treatment groups were then challenged with seven different doses of each of the three parasite strains. After 30 days, the infection status of each offspring *Daphnia* was determined, resulting in data on the fraction of infected *Daphnia* as a function of the exposure dose. The experiment is sketched in Figure 1.

**Figure 1:**
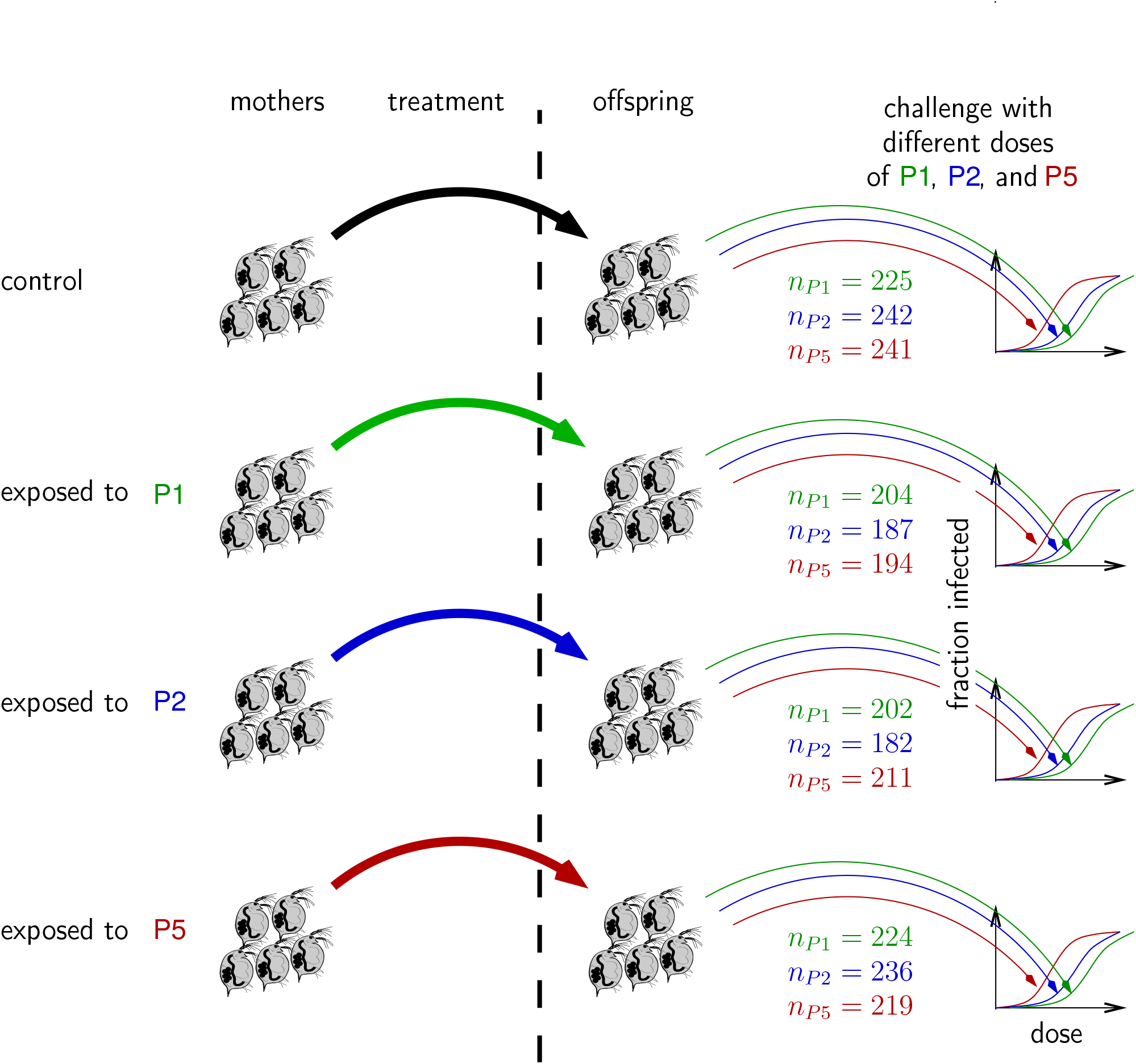
Design of our experiment. Mother *Daphnia* are exposed to three different strain of *Pasteuria ramosa*, P1, P2, or P5. A control cohort of mother *Daphnia* was not exposed. The offspring of these mothers are then exposed to seven different challenge doses of P1, P2, or P5. The sample sizes for each group are indicated on the diagram. They amount to approximately 20–30 individuals per strain and challenge dose. In total we used 2567 individuals.

The infection data for homologous and heterologous challenge groups, as well as the control group are shown in Figure 2. We observed that infection rates increase with increasing dose, and maternal exposure decreases infection rates slightly. Since these data are the result of potentially competing influences of specific and unspecific transgenerational immune priming, a formal method was required to disentangle the effects of maternal exposure on offspring susceptibility.

**Figure 2:**
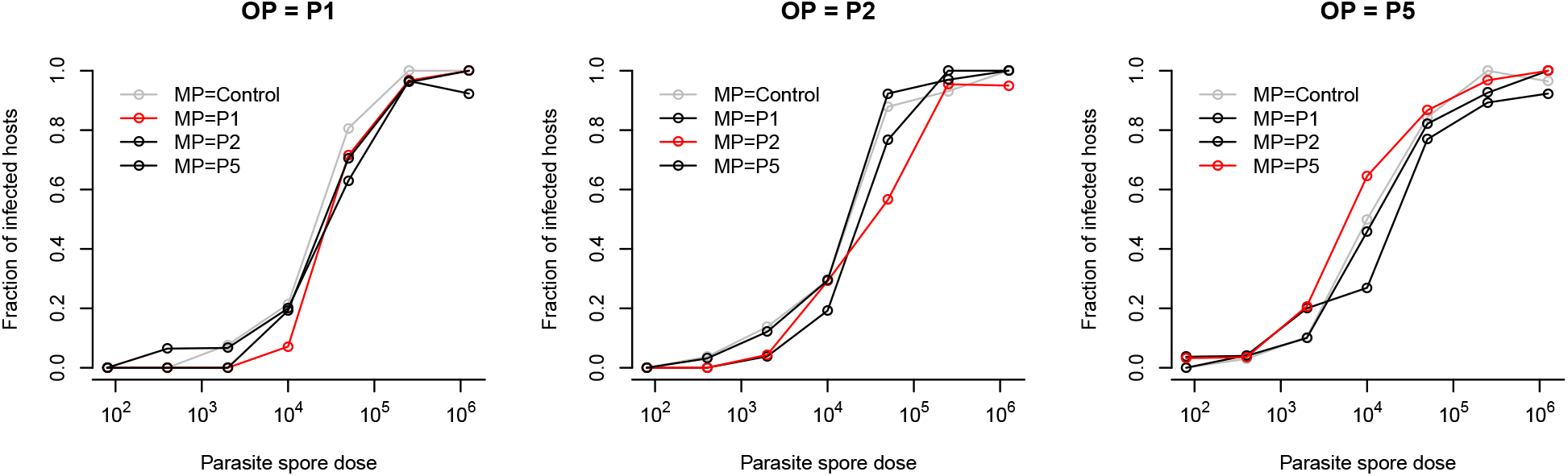
Fraction of infected hosts versus parasite challenge dose for each parasite strain to which offspring *Daphnia* were exposed. OP means “offspring parasite”, the strain to which offspring *Daphnia* were exposed. MP means “mother parasite”, the strain to which mother *Daphnia* were exposed.

### 2.2 Modeling framework

To analyze these data, we extended a mathematical framework developed previously that allowed us to estimate the average infection probability and its inter-individual variance (Regoes et al., 2003, Ben-Ami et al., 2008, 2010). The inspiration for our previous work came from frailty mixing models in mathematical epidemiology (Halloran et al., 1996, Longini and Hallo-ran, 1996, Halloran et al., 2010), but the approaches are also used in microbial risk assessment (Furumoto and Mickey, 1967, Haas, 1999).

Previously, using a single parasite strain, we contrasted exposed mothers to unexposed (control) mothers to determine the potential alterations of susceptibility (Ben-Ami et al., 2008, 2010). We extended this framework by incorporating parameters that capture all conceivable ways of how exposure of the mothers to a specific parasite strain can alter the susceptibility of the offspring to infection. This is needed to analyze our experiments, in which the mother and offspring generation were exposed to three parasite strains in all combinations. Hereby, we separated these potential alterations of susceptibility into two components, a heterologous and a homologous. For example, if the mother *Daphnia* was exposed to P1, its offspring may be less susceptible to challenge with any parasite strain. This would constitute unspecific immune priming against heterologous challenge. Alternatively, maternal exposure to P1 could reduce susceptibility of offspring to P1 specifically, i.e. partially protect against homologous challenge of offspring.

Figure 3 illustrates the decomposition of susceptibility into heterologous and homologous components. The matrix on the left hand side represents the susceptibilities of the offspring of the control group to each parasite strain. The second left panel illustrates unspecific effects of priming with each parasite strain on offspring susceptibility. The third matrix from the right captures a potential memory effect. All of these components are synthesized to the matrix on the left.

**Figure 3:**
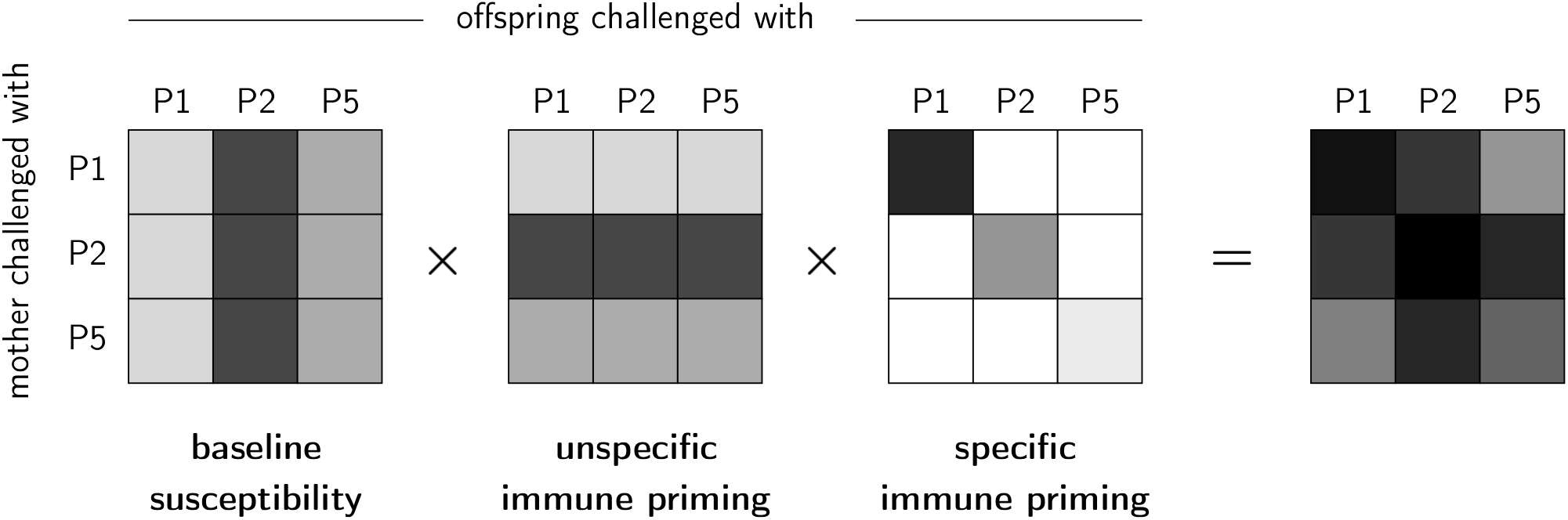
Decomposing the susceptibilities into heterologous and homologous components.

To accomplish this formally, we parameterized the mean susceptibility of *Daphnia b* in the following way: 
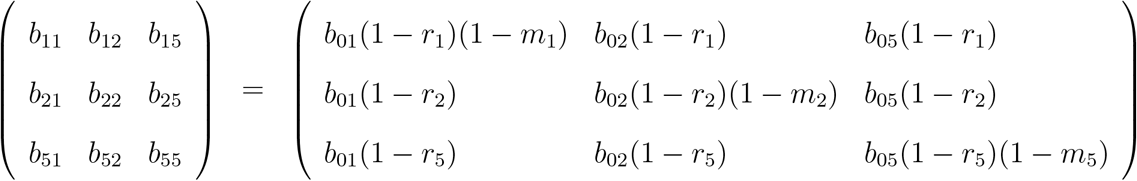

Hereby, *b_ij_* denotes the mean susceptibility to infection per parasite spore

First, we estimated the average susceptibility to each parasite strain *b*_0*j*_ and its variance *v*_0*j*_ from the infection data of control group. For P1, P2, and P5, respectively, we obtain *b*_01_ = 8.73 × 10^−5^, *b*_02_ = 1.68 × 10^−4^, and *b*_05_ = 2.49 × 10^−4^ as the average susceptibilities, and *v*_01_ = 10^−9^, *v*_02_ = 0.73, and *v*_05_ = 0.91 as the susceptibility variances. The model fit to the control data provides the baseline parameters that are needed to determine the quantitative effects of maternal exposure on susceptibility. Figure S1 shows the likelihoods for the control data. Figure 5A shows the fits to the control data.

To study if there is unspecific or specific transgenerational immune priming, we adopt a model selection scheme. Constructing models with increasing complexity and comparing the quality of their fit to the data statistically, we test for immune priming effect. Table 1 lists and defines the models we considered. The simplest model (“0”) is one without any immune priming effects and serves as the null model in our analysis.

**Table 1:** Model variants considered in the model selection scheme.

| Model | Description |
| --- | --- |
| 0 | null model without unspecific or specific immune priming |
| $r$ | unspecific immune priming, equal for each of the three strains |
| $m$ | specific immune priming, equal for each of the three strains |
| $\rho$ | unspecific change in susceptibility variance, equal for each of the three strains |
| $\mu$ | specific change in susceptibility variance, equal for each of the three strains |
| $r-m$ | unspecific and specific immune priming; both effects equal for the three strains |
| $r-\rho$ | unspecific immune priming and unspecific change of the susceptibility variance; both effects equal for the three strains |
| $r-\mu$ | unspecific immune priming and specific change of the susceptibility variance; both effects equal for the three strains |
| $r_i$ | unspecific immune priming; effect may differ between the three strains |
| $r-m_i$ | unspecific and specific immune priming; specific effect may differ between the three strains |
| $r_i-m$ | unspecific and specific immune priming; unspecific effect may differ between the three strains |
| $r-m-\rho$ | unspecific and specific immune priming; unspecific change of susceptibility variance |
| $r-m-\mu$ | unspecific and specific immune priming; specific change of susceptibility variance |
| $r_i-m_i$ | unspecific and specific immune priming; both effects may differ between the three strains |
| $r-m_i-\rho$ | unspecific and specific immune priming; specific effect may differ between the three strains; unspecific change of susceptibility variance |
| $r-m_i-\mu$ | unspecific and specific immune priming; specific effect may differ between the three strains; specific change of susceptibility variance |
| $r-m_i-\rho_i$ | unspecific and specific immune priming; specific effect may differ between the three strains; unspecific change of susceptibility variance that may differ between the three strains |
| $r-m_i-\mu_i$ | unspecific and specific immune priming; specific effect may differ between the three strains; specific change of susceptibility variance that may differ between the three strains |

### 2.3 Evidence for unspecific immune priming

There are four conceivable models that are one step more complex than the null model: “*r*”, “*m*”, “*ρ*”, and “*µ*” (see Table 1 and Figure 4). They allow for unspecific and specific effects on the average susceptibility or its variation. But this potential effect is not allowed to differ between the parasite strain P1, P2, and P5.

**Figure 4:**
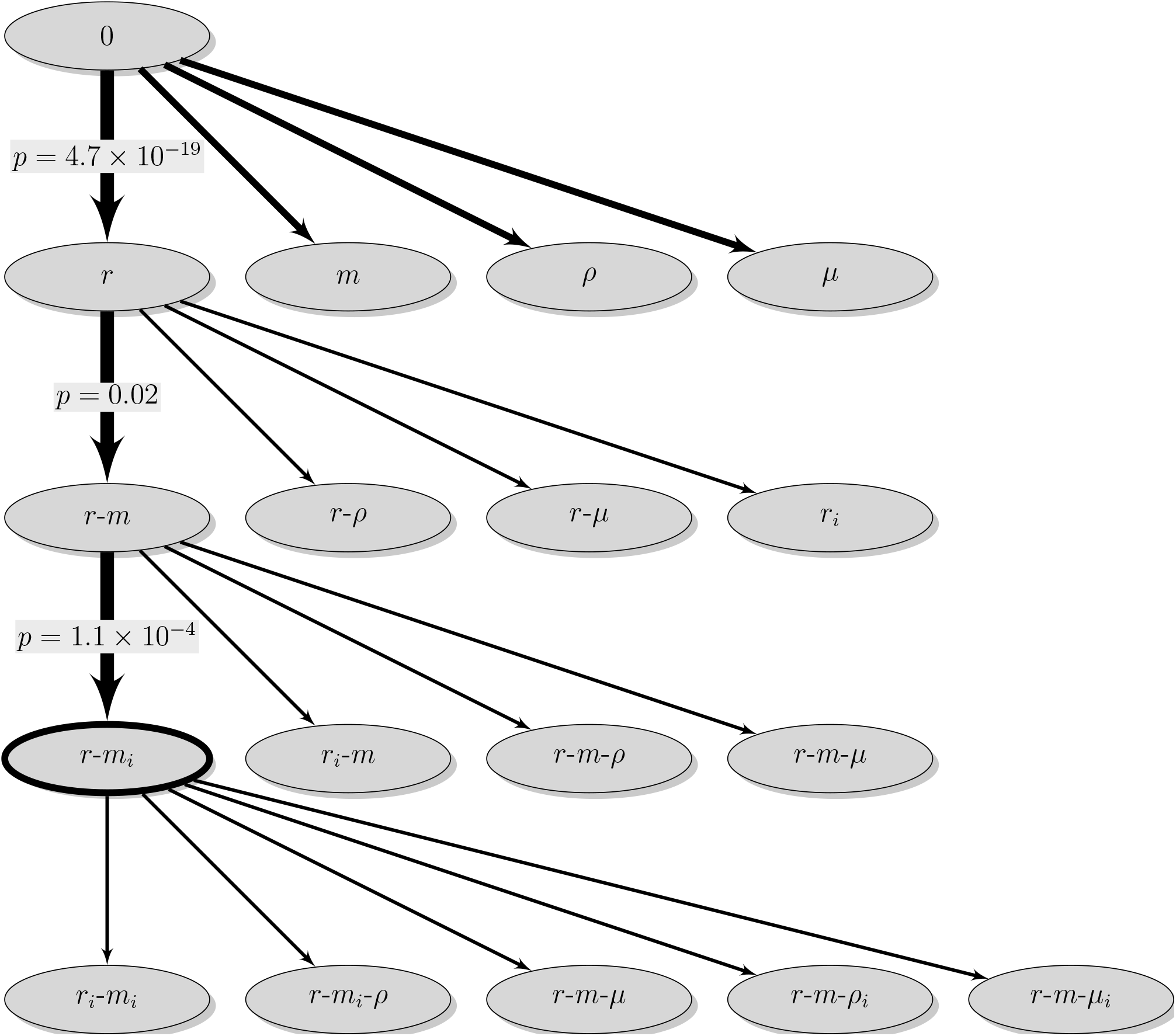
Model selection scheme. The thick lines show the statistically significant model improvements. The thick ellipse circles the most complex model with statistical support.

While all of these model extensions fit significantly better than the null model, the largest improvement in fit arises from the “r” model that describes unspecific, cross-strain immune priming effect (likelihood ratio test: *p* = 4.7 × 10^−19^). We estimate an effect *r* = 0.43. This means that maternal exposure to a parasite reduces the average susceptibility of the offspring to any strain by 43%. Assuming mass-action infection kinetics, this reduction translates into 2.3-fold increase of the ID50.

### 2.4 No evidence for specific immune priming

Because the *r*-model resulted in the largest improvement of model fit we used it as a baseline for the subsequent analysis. We considered four models that are one step more complex than the *r*-model (see Table 1). Biologically the most relevant of these are the *r* − *m*-model, which allows for specific immune priming in addition to the unspecific effect already described in the *r*-model, and the *r_i_*-model that allows the unspecific immune priming effect to differ by parasite strain.

Of the four conceivable models, only the *r* − *m*-model improves the fit significantly (likelihood ratio test: *p* = 0.02; see also Figure 4). Thus, we have evidence for a specific, trans-generational memory of parasite strains. However, the parameter *m* in this model, which describes how well the parasite strain the maternal parasite strain is remembered, is negative: *m* = −0.40. This can be interpreted as specific facilitation of infection, rather than specific protection, and is thus the opposite of immune priming.

### 2.5 Maternal exposure to P5 facilitates infection with P5

The *r* − *m*-model can be extended in various ways (see Table 1 and Figure 4). The most relevant extensions are the *r* − *m_i_*-model and the *r_i_ − m*-model. These two models allow differences in specific and unspecific effect across parasite strains, respectively.

Of all the conceivable extensions, however, only the *r−m_i_*-model improves the fit significantly (likelihood ratio test: *p* = 1.1 × 10^−4^; see also Figure 4). The fit of this model is not improved by any further model extensions (Figure 4 bottom row). Hence, the *r − m_i_*-model represents the model complex enough to capture all aspects related to unspecific and specific immune priming in the data without overfitting them. Figure 5B shows the fits of the best model (*r*-*m_i_* model) to each group of the data.

**Figure 5:**
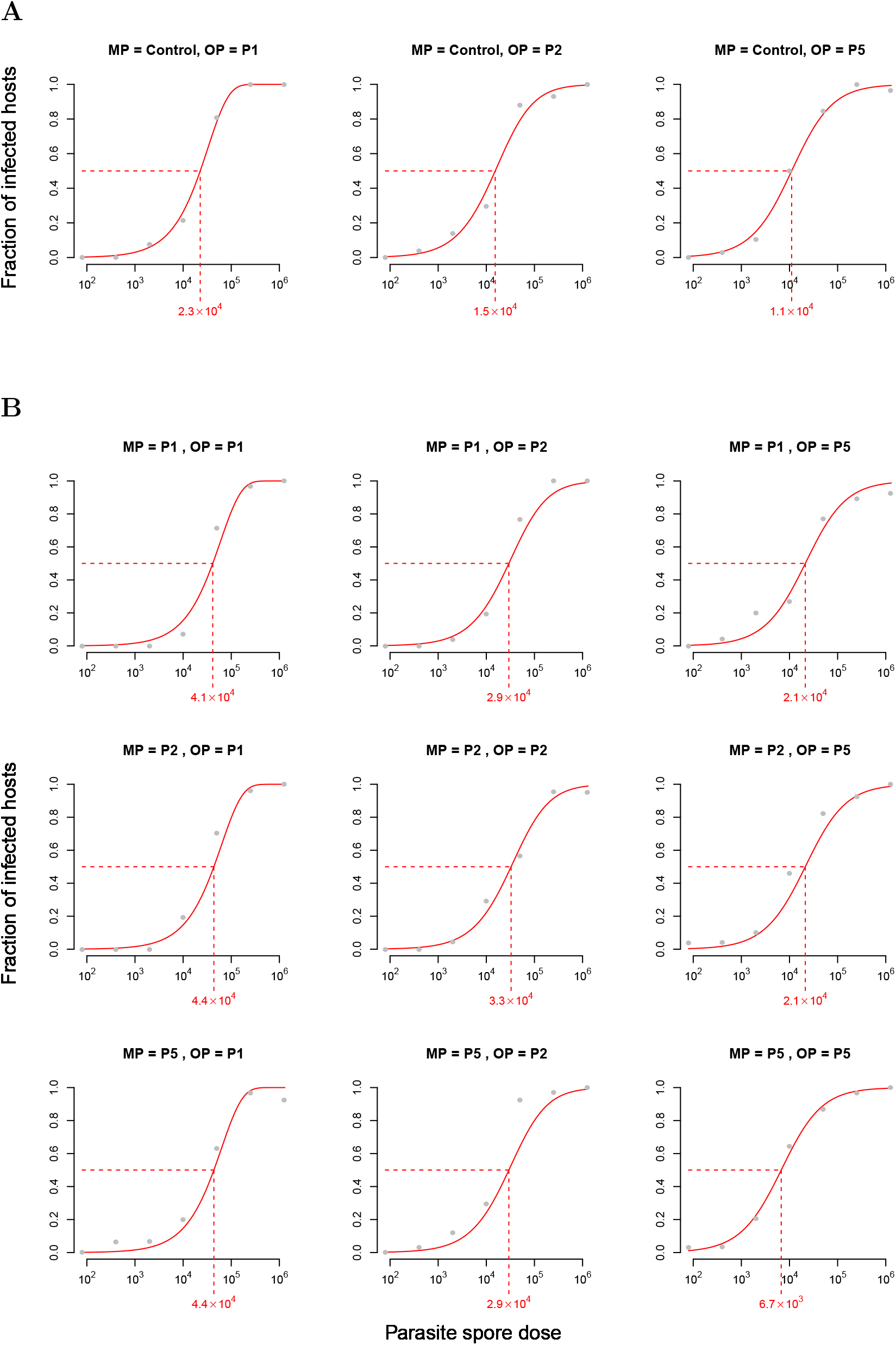
Model fits. (A) Fits of the heterogeneous susceptibility model to the control data. B) Fits of the best model (the *r*-*m_i_* model). The red number below the x-axes gives the spore dose, at which 50% of the hosts are infected (ID50).

The estimate of the unspecific effect of maternal exposure in the *r −m_i_*-model is estimated as *r* = 0.48 with a 95% confidence interval between 0.39 and 0.55. The three parameters describing specific immune priming are estimated as *m*_1_ = −0.054(−0.56, 0.29), *m*_2_ = 0.10(−0.50, 0.46), and *m*_5_ = −2.16(−4.07*, −*0.97). The numbers in brackets give the 95% confidence intervals. Importantly, only *m*_5_ is estimated to be significantly different from 0, and is negative. This means that exposing mothers to P5 facilitates infection with P5.

## 3 Discussion

In the present study, we exposed *Daphnia magna* mothers to three strains of the bacterial parasite *Pasteuria ramosa*, and then exposed the offspring with homologous and heterologous challenges. Our aim was to determine if transgenerational effects of parasite exposure on the distribution of host susceptibility to infection are driven by unspecific or specific immune priming. We found strong evidence of unspecific, cross-strain immune priming, which decreases the susceptibility of host offspring by approximately 50%. We found no evidence of specific immune priming that reduces susceptibility to infection by the same strain the mother had been challenged with (homologous exposure), as compared to other strains (heterologous). However, we found that maternal exposure to one particular parasite strain actually facilitates offspring infection with this parasite, as compared to infections with the other strains.

Our results are different from those of Little et al. (2003). There is no evidence of specific immune priming fine enough to distinguish different strains of *Pasteuria*. It is important to note that we consider a potential effect of previous exposure on mean host susceptibility and its variance, while Little et al. (2003) considered the same effect on infectivity and host fertility. Furthermore, the approximately 1000-fold differences in sensitivities between the two strains used by Little et al. (2003) might indicate a confounding dose effect. Alternatively, specific immune priming might be driven by genotype-by-genotype (GxG) interactions, which are well documented in this system (Carius et al., 2001), insofar the specific host-parasite combinations would be more likely than others to exhibit specific immune priming. The conflict between our study and that of Little et al. (2003) should not be conflated with the more fundamental criticism voiced against studies of invertebrate immunity (Rowley and Powell, 2007, Hauton and Smith, 2007). As many studies of invertebrate immunity, we do not study the molecular basis of the effects we find. But, by relying on the formal concepts and the experimental design principles of population biology, we succeed in elucidating the phenomenological effects of previous exposure on host susceptibility to a surprising level of detail.

The mean susceptibility to isolate P1 and its variance were significantly lower in comparison with P2 and P5. These results are consistent with earlier studies of P1 and two other *Pasteuria* isolates, P3 and P4, which showed that P1 had the lowest infectivity (Ben-Ami et al., 2008). Moreover, in a variety of mixed infections scenarios, P1 was found to be more virulent but produced fewer spores than isolates P3/P4 and clone C1 (obtained from isolate P5, Ben-Ami et al. (2008), Ben-Ami and Routtu (2013)). If virulence is traded off with infectivity, this could influence specific immune priming and transgenerational memory.

Our results are consistent with previous studies of transgenerational effects of *Pasteuria* exposure on *Daphnia* susceptibility (Ben-Ami et al., 2008, 2010). In those studies, we exposed *Daphnia* to one *Pasteuria* isolate. Thus, we could not test for specific immune priming. Some of the parameter estimates we obtained in the present study, however, are inconsistent with those from our previous study (Ben-Ami et al., 2010). Previously, we worked only with P5. We estimated a mean susceptibility of 9.6 × 10^−4^ and 7.2 × 10^−4^ for control and P5-exposed groups, respectively. In contrast, here we get 2.5×10^−4^ and 2.5×10^−4^×(1−0.5)×(1−(−2.2)) = 4×10^−4^.

In other words, in the present study mean susceptibility of the control group was almost fourfold lower than previously, whereas mean susceptibility of the exposed group was less than twofold lower than previously. Consequently, in the present study mean susceptibility of the control group was 33% higher than that of the exposed group, whereas in the previous study it was 38% lower. Although the susceptibility variance of the control group did not change significantly across studies, i.e.,*_v_*_05_ = 0.91 in the present study versus 0.85 previously, for the P5-exposed group we get 1.0, while we got 1.6 previously, a decrease of approximately 38%.

Since the comparison between the present study and the previous one was unplanned, it is not surprising to find differences between the two studies. Nevertheless it is important to carefully consider what factors could have lead to such a divergence between the studies in the mean and variance of offspring susceptibility in the treatments with control and P5-exposed mothers. First, environmental conditions such as food availability and host density can influence maternal effects (Mitchell and Read, 2005, Boots and Roberts, 2012, Michel et al., 2016). The duration of exposure could also influence susceptibility (Bandilla et al., 2005, Wang and Spear, 2016). However, daily food levels of control and exposed treatments and the duration of exposure of exposed treatments in this study were very similar to those in the previous one (Ben-Ami et al., 2010). Second, phenotypic heterogeneity in host susceptibility to environmental and physiological factors, such as molecular differences in immune response (Brites et al., 2008) and within-clone variation in life-history traits (e.g., differences in size at birth; Garbutt and Little (2017)), could also influence the mean and variance of offspring susceptibility. Such heterogeneity would, however, not explain why maternal exposure to parasite isolates P1 or P2 did not facilitate offspring infections as it did for P5. Lastly, the genetic composition of parasite isolate P5 might have changed across studies. Isolates are parasite samples from infected hosts that may contain multiple genotypes (Luijckx et al., 2011). They are a naturally occurring feature of the *Daphnia-Pasteuria* host-parasite system. In the laboratory, isolates are propagated through experimental hosts, to obtain enough spore-carrying cadavers to produce sufficient amounts of spore suspensions. Thus, it might be that over time, some genotypes within the P5 isolate have changed in frequency.

In summary, we present the most extensive data set and analysis of trans-generational immune priming in invertebrates to-date. While we find evidence for specific priming by a specific parasite strain (P5), we do not find any support for the idea of a general specific priming effect (or “memory”) in the *Daphnia-Pasteuria* system.

Formally, our work represents an important contribution not only to the analysis of immune priming effects in ecological systems, but also to the experimental assessment of vaccines. In the epidemiological setting, frailty models have been used to infer the distribution of susceptibilities and vaccine effects (Halloran et al., 2010). In these settings, however, the transmitted dose cannot be controlled, often not even be measured. In experimental settings, dose-response curves are starting to be used for the determination of vaccine effects beyond the average reduction of susceptibility (Langwig et al., 2017). However, an inference framework to investigate specific and unspecific effects of immune priming has not been available to-date. These effects are central to understanding immunity and the effect of vaccines because immune memory and vaccines are typically specific to certain pathogen strains. Because vaccines aim to provide specific protection against certain pathogen strains, our inference framework will therefore be of use much beyond the example of *Daphnia magna* and their parasites that we presented here.

## 4 Materials and Methods

### 4.1 Ethics statement

All applicable institutional and/or national guidelines for the care and use of animals were followed.

### 4.2 Study organisms

*Daphnia magna* Straus is a cyclical parthenogenetic zooplankton, found in a variety of freshwater habitats, such as ponds and rain pools. In nature, many populations are found to be infected by numerous bacterial, microsporidial and fungal parasites (Green, 1974, Ebert, 2005, Goren and Ben-Ami, 2013). In the laboratory, clonal lines can be kept for many generations, allowing to exclude genetic effects experimentally. One of the most common obligate endoparasites of *Daphnia magna* is the bacterium *Pasteuria ramosa* Metchnikoff 1888. This bacterial parasite castrates its host and has a strictly horizontal transmission strategy, by releasing spores from the cadaver of infected *Daphnia* (Ebert et al., 1996, 2016).

### 4.3 Experimental design

Our experimental design is summarized in Figure 1. In brief, we initially either exposed *Daphnia* mothers to one of three *Pasteuria ramosa* isolates (P1: Gaarzerfeld, Germany, 1997; P2: Kaimes, England, 2002; P5: Moscow, Russia), or left them unexposed as controls. These isolates are parasite samples from infected hosts that may contain multiple genotypes (Luijckx et al., 2011). Isolates are a naturally occurring feature of the Daphnia-Pasteuria host-parasite system, and are thus relevant to evolutionary processes in natural populations (Ebert et al., 2016). Despite the potential genetic heterogeneity of the parasite isolates, specific, heritable interactions with the host have been observed (Little et al., 2006).

We subsequently collected the offspring of the mothers, and exposed them to different doses of all three *Pasteuria* isolates, thereby creating one homologous and two heterologous groups per maternal treatment group and dose. We used a single laboratory-maintained *Daphnia magna* clone (HO2 from Hungary) in order to exclude genetic variation among hosts apart from mutations. Isolate P5 was used in a previous study of maternal effects of *Daphnia magna* (Ben-Ami et al., 2010).

For the mothers generation, we placed 4-day-old juveniles individually in 100-mL jars with 20 mL of artificial medium (ADaM; Ebert et al. 1998), and on day 5 all individuals in the exposed treatments were challenged with 50,000 spores of the respective *Pasteuria ramosa* isolate. The animals were fed 1 × 10^6^ algae cells of *Scenedesmus gracilis* per *Daphnia* per day. On day 12, the medium of all animals was replaced with 100 mL of fresh medium, and thereafter medium was replaced every week. The food levels were increased on days 6, 9, 11 and 13 to 2 × 10^6^, 2.5 × 10^6^, 3 × 10^6^ and 8 × 10^6^ algae cells per individual per day, respectively, to accommodate the growing food demand.

For the second generation offspring were collected daily from the experimental mothers and at an age of 4 days were singly placed in 100-mL jars with 20 mL of medium. The offspring of each mother group were randomly assigned to one of seven dose levels (80, 400, 2,000, 10,000, 50,000, 250,000 and 1,250,000 spores/animal) or to a control group. On day 5 all individuals were exposed to their respective parasite strain/dose combination, and after a week the medium of all animals was replaced with 100 mL of fresh medium. Thereafter, medium replacement and feeding schedules were identical to those of the mothers. Both mothers and offspring were kept at 20 *±* 0.5°C, and the light:dark cycle was 16:8. Jars from all treatment groups were distributed randomly across the shelves in a controlled climate room and rearranged frequently to prevent position effects.

Mortality of offspring was recorded daily. Offspring died due to injury inflicted during the separation from their mothers. Only animals that had died after day 16 were checked for disease, because infection cannot be reliably determined earlier. Animals that had died earlier were excluded from the analysis. The experiment ended on day 44, upon which all animals were scored by eye for infection, and their infection status was recorded. When in doubt, we dissected the animal and checked for infection using a phase contrast microscope (300–600X), but we found no discrepancies with our initial diagnosis.

---

**Figure S1:**
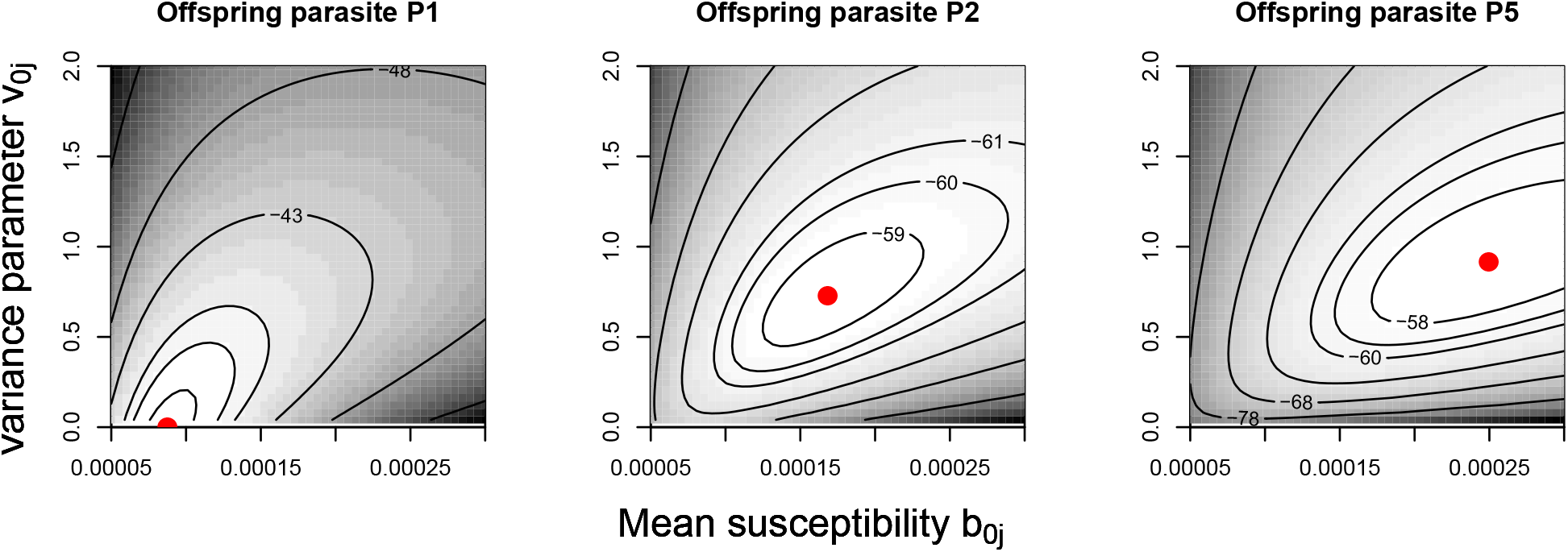
Likelihood surfaces for control data.

